# To dereplicate or not to dereplicate?

**DOI:** 10.1101/848176

**Authors:** Jacob T. Evans, Vincent J. Denef

## Abstract

Our ability to reconstruct genomes from metagenomic datasets has rapidly evolved over the past decade, leading to publications presenting 1,000s, and even more than 100,000 metagenome-assembled genomes (MAGs) from 1,000s of samples. While this wealth of genomic data is critical to expand our understanding of microbial diversity, evolution, and ecology, various issues have been observed in some of these datasets that risk obfuscating scientific inquiry. In this perspective we focus on the issue of identical or highly similar genomes assembled from independent datasets. While obtaining multiple genomic representatives for a species is highly valuable, multiple copies of the same or highly similar genomes complicates downstream analysis. We analyzed data from recent studies to show the levels of redundancy within these datasets, the highly variable performance of commonly used dereplication tools, and to point to existing approaches to account and leverage repeated sampling of the same/similar populations.

---

While initially, the reconstruction of MAGs was only achievable in lower-diversity or highly uneven communities (1), in the past five years reports on the reconstruction of hundreds to thousands of MAGs have become routine (2–5). In the past year, highly automated assembly and binning pipelines have accelerated this trend (6, 7). While these advances open up exciting prospects for addressing questions regarding the physiology, ecology, and evolution of microbial life, MAGs are inherently less reliable than isolate genomes due to their assembly and binning from DNA sequences originating from a mixed community. Various reports have highlighted issues associated with MAGs, including how misassemblies and/or incorrect binning can lead to composite genomes (8, 9) and how fragmented assembly due to strain variation can lead to incomplete genomes that lead to wrong conclusions (10, 11). The latter is a reason why independent assembly of each individual sample is often preferable to avoid assembly fragmentation due to genomic variation between conspecific populations in different samples. However, this often leads to highly similar or identical MAGs being generated across the sample dataset. Multiple tools have been developed to remove redundant MAGs, mainly based on average nucleotide identity between MAGs after sequence alignment using blastn (e.g., pyANI (12)), or faster algorithms combining Mash (13) and gANI (14) or ANIm (15) (e.g., as implemented in dRep (16)).

## Why dereplicate?

Dereplication is the reduction of a set of genomes, typically assembled from metagenomic data, based on high sequence similarity between these genomes. The main reason to do so is that when redundancy in a database of genomes is maintained, the subsequent step of mapping sequencing reads back to this database of genomes leads to sequencing reads having multiple high quality alignments which, depending on the software used and parameters chosen, leads to reads being randomly distributed across the redundant genomes with one random alignment reported from many possible options, or read alignments being reported at all redundant locations. When using these data to make inferences about the relative abundance and population dynamics across samples, relative abundance for the species will look artificially low, and it will appear that multiple ecologically equivalent populations co-occur. Instead, the correct conclusion would be that one more abundant population exists across all samples (Figure 1). This issue has been acknowledged in multiple studies, and authors have chosen varying cutoffs to avoid this issue (e.g., >95% average nucleotide identity (Almeida, 2019); >98% average nucleotide identity (3, 17), >95 % amino acid identity (18), >99.5% amino acid identity (4)).

**Figure 1:**
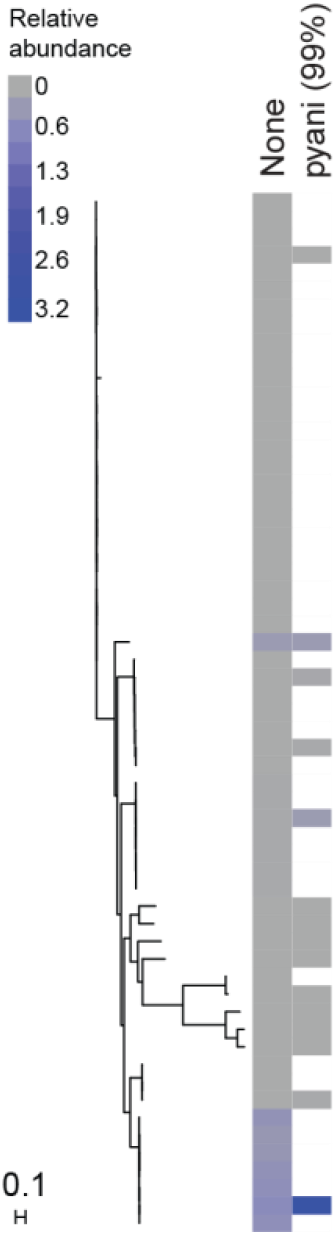
Relative abundance of a set of closely related MAGs from Parks et al, 2017 estimated from read alignment from a metagenomic dataset (SRR1702559) using all genomes in the tree (left column ranging from grey to dark blue) or only those retained after dereplication with pyANI using a 99% ANI cutoff (right column). To calculate relative abundance, all MAGs were combined into a multifasta file. Reads were then mapped to each multifasta fasta file using bwa mem with default parameters (19). Average coverage per contig (grey to blue colors) was computed with pileup.sh from bbtools (https://sourceforge.net/projects/bbmap/). The phylogenetic tree was created by searching for marker genes with pylosift (20) using its default set of marker genes. The genes were then aligned with phylosift, the resulting alignments concatenated, and the tree was created with Fasttree (21) using the -nt and -gtr parameters.

## Why not dereplicate?

Obtaining sequences of multiple individuals of a single population or of individuals of multiple, related populations (a population being defined as individuals of the same species occurring at the same time and place), is valuable as it allows for population genomic analyses that give insights into the intersection between microbial evolution and ecology (22). The standard approach to dereplicate removes genomes based on sequence identity of shared parts of the genome. As such, when removing genomes, in addition to data on single nucleotide polymorphism variation, we may lose information on variability in the auxiliary gene content among representatives from the same species. As an example, we analyzed the effect of dereplication on database auxiliary gene content using two of the most commonly used tools (dRep and redundancy removal based on pyANI results). We used a set of 46 *Microcystis aeruginosa* MAGs we previously generated with extensive manual curation (11). The ANI between pairs of these 46 genomes averages 96.4%. Out of a total of 9,175 unique gene clusters across the 46 MAGs, dereplication led to the removal of up to 2,228 auxiliary genes when using dRep gANI with a 96.5 % cutoff (used for species delineation using genome sequences (14)) (Fig. 2). On the other extreme, using dRep default, no genomes were removed from the MAG set thus no gene clusters were lost, while intermediate numbers of gene clusters were removed when using pyANI (213) and dRep gANI (447) at 99% thresholds.

**Figure 2:**
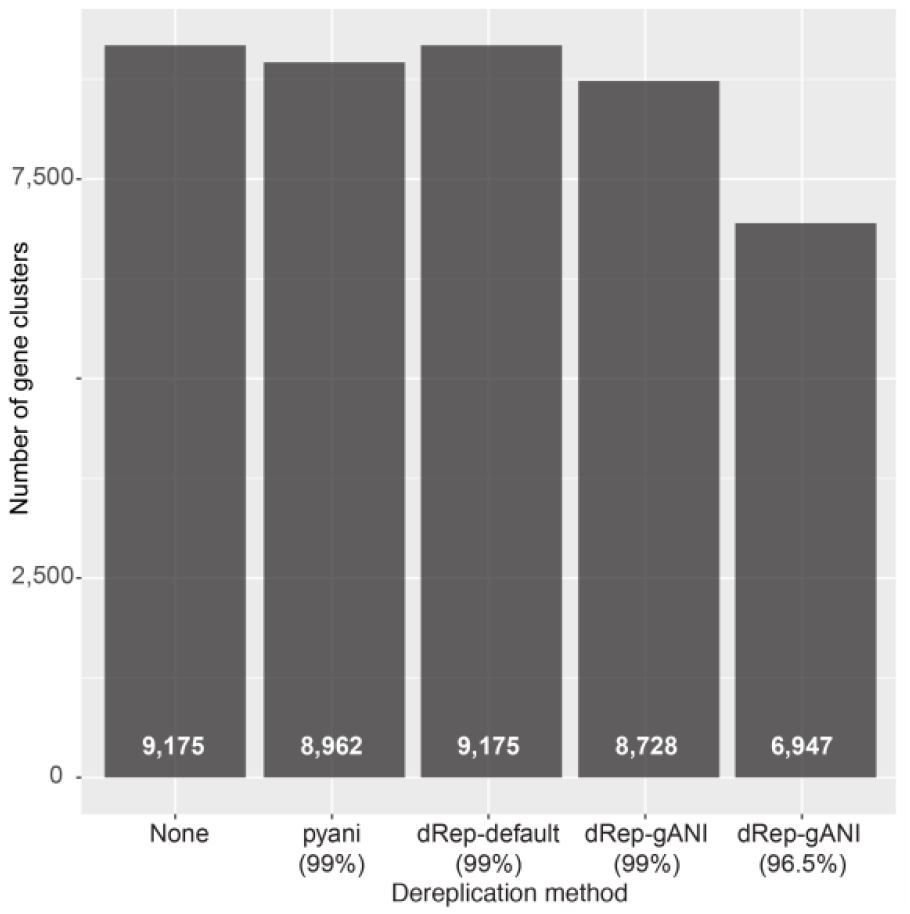
Retained gene clusters of the Microcystis pangenome when using different dereplication tools and settings.

## Variable performance of commonly used software

As already indicated from the analysis in Figure 2, different dereplication tools lead to different outcomes, even when using the same sequence identity cutoffs. Using publicly available MAG data sets (4, 6, 7), we evaluated the performance of two commonly used dereplication tools, dRep and pyANI. For dRep, we used the default parameters based on genome-wide alignments using animf with nucmer and a cutoff of 99% (23), and dRep using the gANI option that does gene-based alignments using nucmer with a cutoff of 99% and 96.5%. For pyANI, we used a 99% ANI cutoff and sequence identity is calculated using blast-based genome-wide alignments. While slower, we consider it the reference to compare to due to the higher accuracy of blast-based alignments (24). Prior to running pyANI, in order to avoid calculating ANI for distantly related pairwise comparisons so as to reduce computation time, groups of MAGs were formed by calculating pairwise distances using Mash (default parameters; (13)). The computed pairwise distances were then used to cluster genomes into similar groups with hierarchical clustering using a custom python script with fcluster from SciPy (http://www.scipy.org/) with a threshold of 2. pyANI was then run within each group created from the clustering.

First, we performed a comprehensive analysis of a set of 7,800 genomes generated from 1,550 public metagenomes (4). In this study, no dereplication was done for most analyses except for building the tree represented in Figure 2 in this study. For the latter analysis, dereplication was performed by removing genomes with an amino-acid identity (AAI) ≥99.5% as calculated using CompareM (https://github.com/dparks1134/CompareM), resulting in the removal of 27.5% of all MAGs. In our own analyses, relative to the pyANI reference (32.9% removal), default dRep removed fewer genomes (19.3%), while the gANI dRep approach removed more MAGs (48.1% (99% ANI), 56.9% (96.5% ANI)) (Fig. 3A). A closer look at one cluster of related MAGs indicated that dRep gANI regularly removed genomes that did not require removal, while dRep with default parameters was not removing a sufficient number of MAGs (Fig. 3D).

**Figure 3:**
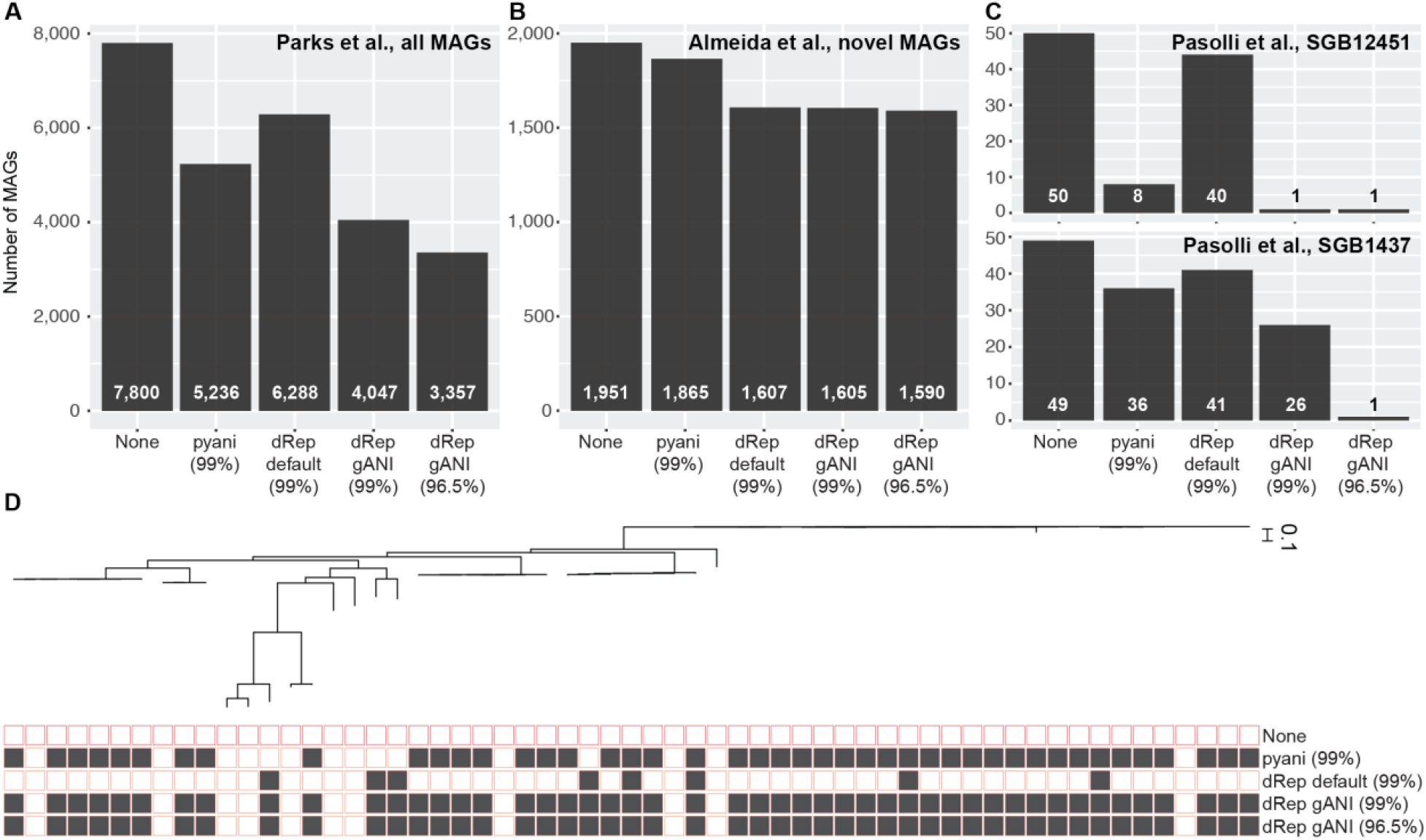
Tool-dependent effects of dereplication. (A-C) Number of MAGs remaining after dereplication tools were run. (D) Phylogenetic tree of the same group of MAGs used in Figure 1, showing differential removal of MAGs after dereplication (filled square indicates MAG was removed after dereplication). dRep default does not remove multiple near-identical MAGs, while dRep-gANI removes MAGs that are more distantly related than the 99% or 96.5% ANI cutoff.

For a recent study that generated more than 90,000 MAGs (6), we performed our comparative dereplication analysis on the 1,952 uncultured bacteria species that were identified and focused by the authors. These were MAGs not classified at the species level in current databases that had been dereplicated by removing less complete MAGs that shared ANI > 95% across 60% of their sequence length. In this case, pyANI removed four times fewer MAGs than the different implementations of dRep (Fig. 3B). In contrast with our preceding analyses, dRep default removed more MAGs than pyANI, potentially due to the fact that the authors had already derpelicated their MAG set at 95% ANI. Finally, we analyzed two MAG groups, clustered at the species level (95% ANI) by the authors of a recent study generating more than 150,000 MAGs (7). In this case, dRep-default again removed fewer MAGs than pyANI, while dRep using gANI removed many more MAGs (Fig. 3C).

## Available approaches to leverage sampling of between-population variation

Several tools have been developed to maintain the auxiliary genomes of closely related strains while avoiding redundancy when tracking strain-resolved population dynamics in the environment using metagenomic data (reviewed in (11)). They typically use metagenomic data in combination with a genomic database of genomes of closely related isolates or MAGs based on whether alleles of shared genes (StrainPhlAn (25); ConStrains (26)), strain-specific auxiliary genes (PanPhlAn (27))}, or both are present in a sample (MIDAS (28)). Similarly, the Anvi’o package incorporates a metapangenome workflow that reduces a set of user-defined conspecific genomes to gene clusters representing core and auxiliary genes and then estimates strain abundances across metagenomic datasets (29). In principle, all of these approaches avoid the issues associated with database redundancy highlighted in Fig. 1, and loss of population-specific auxiliary genes highlighted in Fig. 2. Although variant identification errors do remain, which are tool and likely database and metagenomic dataset dependent, this has been reported to be as low as 0.1% (25). While potential issues with these approaches have not been fully evaluated, analyses focusing on populations where the dominant strain can be more readily resolved have been able to go as far as tracking *in situ* bacterial evolution in environmental biofilms and the human gut (30, 31).

## Conclusions

Genome-centric metagenomics has opened a view onto the undescribed branches of the tree of life (32). Yet, full awareness of the risks associated with MAGs is needed to avoid misinterpretation of the data and populating databases with questionable genomes. Dereplication is a step carried out by many researchers as part of metagenomic informatic pipelines, but we highlight large differences between commonly used tools in how many genomes are removed. Tools able to resolve closely related genomes exist and may circumvent issues with redundancy while maximally leveraging all data contained in MAGs from conspecific population. As the ability to resolve closely related genomes is dependent on the genetic distance between genomes in the database and between database genomes and those of sampled populations, these tools need broader adaptation and evaluation to fully evaluate their accuracy. This in turn may lead to guidelines for a minimum level of dereplication necessary to enable their use.

## Code availability

All code written and used for the analyses described in this manuscript can be found at https://github.com/DenefLab/Dereplication-Letter-Code.

## Acknowledgments

This research was supported by funding from the National Science Foundation to VJD (NSF EAGER 1737680).

## References

1. Tyson GW, Chapman J, P. H, Allen EE, Ram RJ, Richardson PM, Solovyev VV, Rubin EM, Rokhsar DS, Banfield JF. 2004. Community structure and metabolism through reconstruction of microbial genomes from the environment. Nature 428:37–43.

2. Brown CT, Hug LA, Thomas BC, Sharon I, Castelle CJ, Singh A, Wilkins MJ, Wrighton KC, Williams KH, Banfield JF. 2015. Unusual biology across a group comprising more than 15% of domain Bacteria. Nature 523:208.

3. Anantharaman K, Brown CT, Hug LA, Sharon I, Castelle CJ, Probst AJ, Thomas BC, Singh A, Wilkins MJ, Karaoz U. 2016. Thousands of microbial genomes shed light on interconnected biogeochemical processes in an aquifer system. Nature communications 7:13219.

4. Parks DH, Rinke C, Chuvochina M, Chaumeil P-A, Woodcroft BJ, Evans PN, Hugenholtz P, Tyson GW. 2017. Recovery of nearly 8,000 metagenome-assembled genomes substantially expands the tree of life. Nature microbiology 2:1533.

5. Stewart RD, Auffret MD, Warr A, Wiser AH, Press MO, Langford KW, Liachko I, Snelling TJ, Dewhurst RJ, Walker AW. 2018. Assembly of 913 microbial genomes from metagenomic sequencing of the cow rumen. Nature communications 9:870.

6. Almeida A, Mitchell AL, Boland M, Forster SC, Gloor GB, Tarkowska A, Lawley TD, Finn RD. 2019. A new genomic blueprint of the human gut microbiota. Nature 568:499.

7. Pasolli E, Asnicar F, Manara S, Zolfo M, Karcher N, Armanini F, Beghini F, Manghi P, Tett A, Ghensi P. 2019. Extensive unexplored human microbiome diversity revealed by over 150,000 genomes from metagenomes spanning age, geography, and lifestyle. Cell 176:649–662. e620.

8. Koutsovoulos G, Kumar S, Laetsch DR, Stevens L, Daub J, Conlon C, Maroon H, Thomas F, Aboobaker AA, Blaxter M. 2016. No evidence for extensive horizontal gene transfer in the genome of the tardigrade Hypsibius dujardini. Proceedings of the National Academy of Sciences 113:5053–5058.

9. Shaiber A, Eren AM. 2019. Composite Metagenome-Assembled Genomes Reduce the Quality of Public Genome Repositories. mBio 10:e00725–00719.

10. Hug LA, Thomas BC, Sharon I, Brown CT, Sharma R, Hettich RL, Wilkins MJ, Williams KH, Singh A, Banfield JF. 2016. Critical biogeochemical functions in the subsurface are associated with bacteria from new phyla and little studied lineages. Environ Microbiol18:159–173.

11. Jackrel SL, White JD, Evans JT, Buffin K, Hayden K, Sarnelle O, Denef VJ. 2019. Genome evolution and host-microbiome shifts correspond with intraspecific niche divergence within harmful algal bloom-forming Microcystis aeruginosa. Mol Ecol 28:3994–4011.

12. Pritchard L, Glover RH, Humphris S, Elphinstone JG, Toth IK. 2016. Genomics and taxonomy in diagnostics for food security: soft-rotting enterobacterial plant pathogens. Analytical Methods 8:12–24.

13. Ondov BD, Treangen TJ, Melsted P, Mallonee AB, Bergman NH, Koren S, Phillippy AM. 2016. Mash: fast genome and metagenome distance estimation using MinHash. Genome Biol 17:132.

14. Varghese NJ, Mukherjee S, Ivanova N, Konstantinidis KT, Mavrommatis K, Kyrpides NC, Pati A. 2015. Microbial species delineation using whole genome sequences. Nucleic Acids Res 43:6761–6771.

15. Richter M, Rosselló-Móra R. 2009. Shifting the genomic gold standard for the prokaryotic species definition. Proceedings of the National Academy of Sciences 106:19126–19131.

16. Olm MR, Brown CT, Brooks B, Banfield JF. 2017. dRep: a tool for fast and accurate genomic comparisons that enables improved genome recovery from metagenomes through de-replication. ISME J 11:2864.

17. Solden LM, Naas AE, Roux S, Daly RA, Collins WB, Nicora CD, Purvine SO, Hoyt DW, Schückel J, Jørgensen B. 2018. Interspecies cross-feeding orchestrates carbon degradation in the rumen ecosystem. Nature Microbiology 3:1274.

18. Woodcroft BJ, Singleton CM, Boyd JA, Evans PN, Emerson JB, Zayed AA, Hoelzle RD, Lamberton TO, McCalley CK, Hodgkins SB. 2018. Genome-centric view of carbon processing in thawing permafrost. Nature 560:49.

19. Li H. 2013. Aligning sequence reads, clone sequences and assembly contigs with BWA-MEM. arXiv preprint arXiv:13033997.

20. Darling AE, Jospin G, Lowe E, Matsen IV FA, Bik HM, Eisen JA. 2014. PhyloSift: phylogenetic analysis of genomes and metagenomes. PeerJ 2:e243.

21. Price MN, Dehal PS, Arkin AP. 2009. FastTree: Computing Large Minimum Evolution Trees with Profiles instead of a Distance Matrix. Mol Biol Evol 26:1641–1650.

22. Polz MF, Rajora OP. 2019. Population Genomics: Microorganisms. Springer.

23. Kurtz S, Phillippy A, Delcher AL, Smoot M, Shumway M, Antonescu C, Salzberg SL. 2004. Versatile and open software for comparing large genomes. Genome Biol 5:R12.

24. Yoon S-H, Ha S-m, Lim J, Kwon S, Chun J. 2017. A large-scale evaluation of algorithms to calculate average nucleotide identity. Antonie Van Leeuwenhoek 110:1281–1286.

25. Truong DT, Tett A, Pasolli E, Huttenhower C, Segata N. 2017. Microbial strain-level population structure and genetic diversity from metagenomes. Genome Res 27:626–638.

26. Li D, Liu C-M, Luo R, Sadakane K, Lam T-W. 2015. MEGAHIT: an ultra-fast single-node solution for large and complex metagenomics assembly via succinct de Bruijn graph. Bioinformatics 31:1674–1676.

27. Scholz M, Ward DV, Pasolli E, Tolio T, Zolfo M, Asnicar F, Truong DT, Tett A, Morrow AL, Segata N. 2016. Strain-level microbial epidemiology and population genomics from shotgun metagenomics. Nature methods 13:435.

28. Nayfach S, Rodriguez-Mueller B, Garud N, Pollard KS. 2016. An integrated metagenomics pipeline for strain profiling reveals novel patterns of bacterial transmission and biogeography. Genome Res 26:1612–1625.

29. Delmont TO, Eren AM. 2018. Linking pangenomes and metagenomes: the Prochlorococcus metapangenome. PeerJ 6:e4320.

30. Denef VJ, Banfield JF. 2012. In situ evolutionary rate measurements show ecological success of recently emerged bacterial hybrids. Science 336:462–466.

31. Garud NR, Good BH, Hallatschek O, Pollard KS. 2019. Evolutionary dynamics of bacteria in the gut microbiome within and across hosts. PLoS Biol 17:e3000102.

32. Hug LA, Baker BJ, Anantharaman K, Brown CT, Probst AJ, Castelle CJ, Butterfield CN, Hernsdorf AW, Amano Y, Ise K. 2016. A new view of the tree of life. Nature microbiology 1:16048.

